# The genome sequence of the scarce swallowtail, *Iphiclides podalirius*

**DOI:** 10.1101/2022.03.31.486480

**Authors:** Alexander Mackintosh, Dominik R. Laetsch, Tobias Baril, Sam Ebdon, Paul Jay, Roger Vila, Alex Hayward, Konrad Lohse

**Affiliations:** Institute of Evolutionary Biology, University of Edinburgh, Edinburgh, EH9 3FL, UK; Centre for Ecology and Conservation, University of Exeter, Penryn Campus, Cornwall, TR10 9FE, UK; Ecologie Systématique Evolution, Bâtiment 360, CNRS, AgroParisTech, Université Paris-Saclay, 91400 Orsay, France; Institut de Biologia Evolutiva (CSIC - Universitat Pompeu Fabra), Passeig Marítim de la Barceloneta 37, ESP-08003 Barcelona, Spain

## Abstract

The scarce swallowtail, *Iphiclides podalirius* (Linnaeus, 1758), is a species of butterfly in the family Papilionidae. Here we present a chromosome-level genome assembly for *I. podalirius* as well as gene and transposable element annotations. We estimate heterozygosity within different partitions of the genome and find a negative correlation between chromosome length and heterozygosity at fourfold-degenerate sites. This high quality genome assembly, the first for any species in the tribe Leptocircini, will be a valuable resource for population genomics in the genus *Iphiclides* and comparative genomics more generally.

## 1 Introduction

The scarce swallowtail, *Iphiclides podalirius* (Linnaeus, 1758), is a widespread butterfly species in the family Papilionidae. The species is common in open habitats in the Palearctic, ranging from France to Western China, but is absent from Northern areas (e.g. Scandinavia and the British Isles) and some Mediterranean Islands (e.g. Sardinia, where only occasional records exist). *Iphiclides podalirius* is generally bivoltine, the larvae feed mainly on different species of *Prunus*, principally *P. spinosa*, and overwinter in the pupal stage.

The genus *Iphiclides* belongs to the tribe Leptocircini (kite swallowtails), which diverged from other taxa in the subfamily Papilioninae about 55 MYA ago (Allio *et al*. 2019), and only includes two other species: *I. podalirinus*, which is restricted to Central Asia and *I. feisthamelii*, the sister taxon of *I. podalirius*, which lives in Northern Africa and the Iberian Peninsula. The controversy about the taxonomic status of *I. feisthamelii*, which has been regarded as a subspecies of *I. podalirius* (Godart and Duponchel 1832; Tolman and Lewington 2009; Wiemers and Gottsberger 2010), has only recently been resolved; although the two species have no known differences in ecology or life history and share mitochondrial haplotypes (Dincă *et al*. 2015), Gaunet *et al*. (2019) show that they differ consistently in wing patterns (including UV reflectance of males), genital morphology and nuclear DNA and are separated by a narrow hybrid zone in Southern France (Descimon and Mallet 2009).

Although Allio *et al*. (2019) have previously generated Illumina shotgun data for *I. podalirius*, the assemblies based on these data (Allio *et al*. 2019; Ellis *et al*. 2021) are highly fragmented (with an N50 of 0.6 and 1.7 Kb respectively). More generally, while chromosome level assemblies are available for several swallowtail butterflies in the genus *Papilio* (Lu *et al*. 2019), similarly contiguous genome assemblies are lacking for other tribes within Papilioninae.

Here we present a chromosome-level genome assembly for *Iphiclides podalirius*, as well as gene and transposable element annotations. We use this assembly to investigate how heterozygosity in the reference individual varies both between genomic partitions and chromosomes.

## 2 Materials and methods

### 2.1 Sampling

Two male individuals (MO IP 500 and MO IP 504) were collected with a hand net at Le Moulin de Bertrand, Saint-Martin-de-Londres, Montpellier, France and flash frozen in liquid nitrogen. High-molecular-weight (HMW) DNA was extracted from the thorax of one of these individuals (MO IP 504) using a salting out extraction protocol as described in Mackintosh *et al*. (2021).

### 2.2 Sequencing

A SMRTbell sequencing library was generated from the HMW extraction of MO IP 504 by the Exeter Sequencing Service. This was sequenced on three SMRT cells on a Sequel I instrument to generate 24.0 Gb of Pacbio continuous long read (CLR) data.

A chromium 10X library was prepared from the HMW extraction at the Cancer Research UK Cambridge Institute, UK. This library was sequenced by Edinburgh Genomics (EG) on a single HiSeqX lane, generating 25.3 Gb of paired-end reads after processing with Long Ranger v2.2.2 (Marks *et al*. 2019). However, the weighted mean molecule size of these data (12.86 kb) limited its use for scaffolding of the Pacbio assembly and the reads were therefore only used for polishing. Hereafter this data is simply referred to as Illumina WGS.

In addition, tissue from MO IP 500 was used for chromatin conformation capture (HiC) sequencing. The HiC reaction was performed using an Arima-HiC kit, following the manufacturer’s instructions for flash frozen animal tissue. An NEBNext Ultra II library, prepared by EG, was sequenced on an Illumina MiSeq, generating 4.7 Gb of paired-end reads.

Paired-end RNA-seq data were generated for individual MO IP 504. For details on RNA extraction see Ebdon *et al*. (2021).

### 2.3 Genome assembly

Pacbio reads *≥* 2kb (40.4x coverage) were assembled using wtdbg2.5 (Ruan and Li 2020) with the options: -L 2000 -x sq -g 400m. Errors in the consensus sequence were corrected by three rounds of Pacbio CLR polishing and two rounds of Illumina WGS polishing using Racon v1.4.10 (Vaser *et al*. 2017) while retaining any unpolished sequences. Contigs belonging to non-target organisms were identified using blobtools v1.1.1(Laetsch and Blaxter 2017) and subsequently removed. Finally, duplicated regions and contigs with extremely low (<5x) or high (>200x) coverage were identified and removed with purge dups v1.2.5 (Guan *et al*. 2020). Mapping of long-reads and short-reads for the above steps were performed with minimap2 v2.17 and bwa-mem v0.7.17 respectively (Li 2018, 2013).

The HiC and RNA-seq reads were adapter and quality trimmed with fastp v0.2.1 (Chen *et al*. 2018). The trimmed HiC reads were aligned to the contig-level assembly with Juicer v1.6 (Durand *et al*. 2016).Scaffolding was performed with 3d-dna v180922 using default parameters (Dudchenko *et al*. 2017). The scaffolded assembly and HiC map generated by 3d-dna was visualised and manually curated in Juicebox v1.11.08 (Robinson *et al*. 2018).

Gene completeness was evaluated using BUSCO v5.0.0 with the insecta odb10 dataset (n=1367) (Manni *et al*. 2021). Kmer QV was calculated using Merqury v1.1 (Rhie *et al*. 2020).

The mitochondrial genome was assembled and annotated using the Mitofinder pipeline v1.4 (Allio *et al*. 2020). Illumina WGS reads were assembled with metaSPAdes v3.14.1 (Nurk *et al*. 2017) and tRNAs were annotated with MiTFi (Juühling *et al*. 2011).

### 2.4 Genome annotation

Transposable elements (TEs) were annotated using the Earl Grey TE annotation pipeline (https://github.com/TobyBaril/EarlGrey, Baril *et al*. 2021; Platt *et al*. 2016; Wong and Simakov 2018; Smit *et al*. 2015; Xu and Wang 2007; Ou and Jiang 2019; Rubino and Creevey 2014; Jurka *et al*. 2005; Hubley *et al*. 2015; Flynn *et al*. 2020) as in Mackintosh *et al*. (2021).

Following gene annotation, 5’ and 3’ gene flanks were defined as those that were 20kb upstream and downstream of genes. We define regions as intergenic space if they are neither genic (start/stop codons, exons, and introns) nor gene flanks. Bedtools intersect v2.27.1 (Quinlan and Hall 2010) was used to determine overlap (-wao) between TEs and genomic features. Following this, quantification and plotting was performed in R, using the tidyverse package (Wickham *et al*. 2019; R Core Team 2021; RStudio Team 2020).

The trimmed RNA-seq reads were mapped to the assembly with HISAT2 v2.1.0 (Kim *et al*. 2019). The softmasked assembly and RNA-seq alignments were used for gene prediction with braker2.1.5 (Hoff *et al*. 2015, 2019; Li *et al*. 2009; Barnett *et al*. 2011; Lomsadze *et al*. 2014; Buchfink *et al*. 2015; Stanke *et al*. 2006, 2008). Gene annotation statistics were calculated with GenomeTools v1.6.1 (Gremme *et al*. 2013).

Functional annotation was carried out using InterProScan v5.50-84.0 (Jones et al. 2014) and the Pfam-33.1 database (Mistry *et al*. 2020).

### 2.5 Estimating heterozygosity

Heterozygosity was estimated within different partitions of the genome by first mapping the Illumina WGS reads to the assembly with bwa-mem, marking duplicated alignments with sambamba 0.8.1 (Tarasov *et al*. 2015), and calling variants with freebayes v1.3.2-dirty (Garrison and Marth 2012). Variants were normalised with bcftools v1.8 (Danecek *et al*. 2021), decomposed with vcfallelicprimitives (Garrison *et al*. 2021), filtered (QUAL*>* 1) and subset to biallelic SNPs with bcftools. Callable sites were delimited with mosdepth v0.3.2 (Pedersen and Quinlan 2017), by applying a coverage threshold between 8 and 95 (inclusive). Callable sites were further restricted by removing all sites which failed VCF filters. Fourfold-degenerate (4D) and zerofold-degenerate (0D) sites were identified using partition cds.py v0.2 (see Data availability), based on the CDS BED file (where the 4th column contains transcript ID), the genome FASTA and the VCF file. Bedtools v2.30.0 was used to intersect callable regions of the genome with intergenic, intronic, exonic, 4D, and 0D sites. Heterozygosity — the number of heterozygous bi-allelic SNPs divided by the number of callable sites — and callable sites of each genomic partition are listed in Table 1.

**Table 1:** Estimates of heterozygosity in different partitions of the genome.

| Partition | Callable sites (Mb) | Heterozygosity |
| --- | --- | --- |
| All | 420.3 | 0.00598 |
| Exonic | 19.3 | 0.00295 |
| Intronic | 126.1 | 0.00615 |
| Intergenic | 275.1 | 0.00609 |
| 0D | 12.4 | 0.00157 |
| 4D | 3.1 | 0.00680 |

## 3 Results and Discussion

### 3.1 Genome assembly

We sequenced the genome of a male individual (Figure 1A and 1B) by generating both Pacbio continuous long-reads (55.7-fold coverage) and Illumina paired-end short-reads (58.8-fold coverage). From these data, we assembled a haploid representation of the genome consisting of 265 contigs with a total span of 430.6 Mb. The span of the contig assembly is similar to both a genome size estimate based on flow cytometry of male individuals (386.6 Mb, Mackintosh *et al*. 2019) and the size of a previous assembly for this species based only on Illumina data (390.9 Mb, Allio *et al*. 2019). We scaffolded the contigs with Arima HiC data (11.0-fold coverage) generated from a different male individual collected at the same locality (Figure 1C and 1D). Scaffolding generated 30 chromosome-scale sequences which together account for 99.5% of the total assembly length and range from 6.8 to 21.1Mb in size (Figure S1). The assembly has a contig and scaffold N50 of 5.2 Mb and 15.1 Mb, respectively.

**Figure 1:**
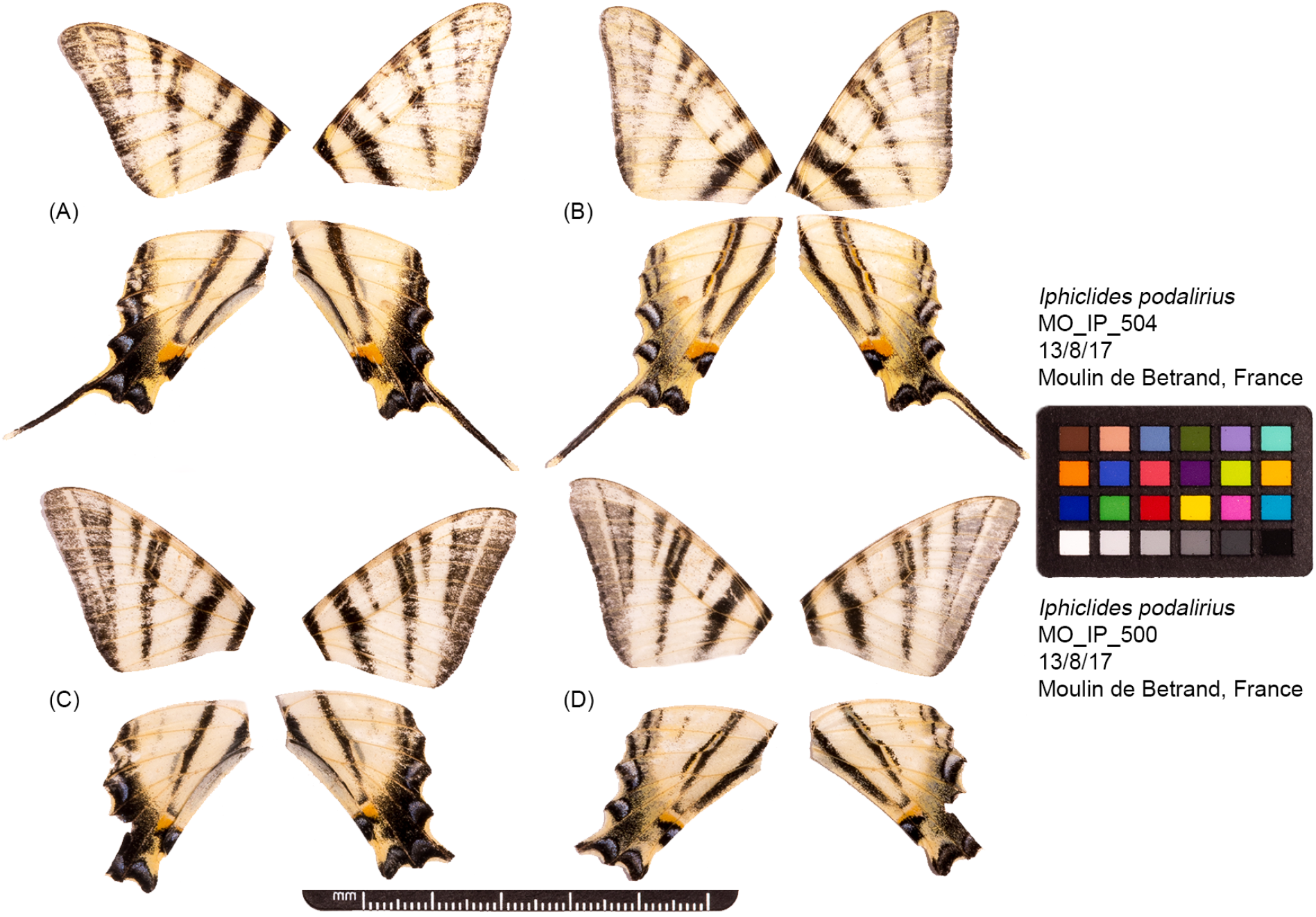
Fore and hind wings of the two *I. podalirius* individuals used to generated the genome sequence. (A) Dorsal and (B) ventral surface view of wings of specimen MO IP 504, used to generate Pacbio long-read, Illumina WGS short-read, and Illumina RNA-seq short-read data. (C) Dorsal and (D) ventral surface view of wings of specimen MO IP 500, used to generate HiC data.

The BUSCO analysis shows that the assembly contains the majority of expected single copy orthologues with little duplication (S:96.8%, D:0.2%, F:0.6%, M:2.4%). The Phred quality of the consensus sequence, estimated using solid kmers in the short-read data, is 35.8.

We assembled a circularised mitochondrial genome of 15,396 bases, containing 13 protein coding genes, 22 tRNA genes, and two rRNA genes. The sequence can be aligned co-linearly with the mitochondrial genome of *Graphium doson* (Kong *et al*. 2019), another species in the tribe Leptocircini, demonstrating that these mitochondrial genomes have not undergone any rearrangements.

### 3.2 Genome annotation

Transposable elements (TEs) compromise 32.81% of the genome assembly (Table S2, Figure 2A). The assembly contains representation from all major TE types (Table S2): the most abundant TEs are long interspersed nuclear elements (LINEs), which constitute 11.01% of the assembly and 33.56% of total TE sequence. Recent activity is high in LINEs and there is also evidence for a very recent increase in LTR element abundance (Figure 2B).

**Figure 2:**
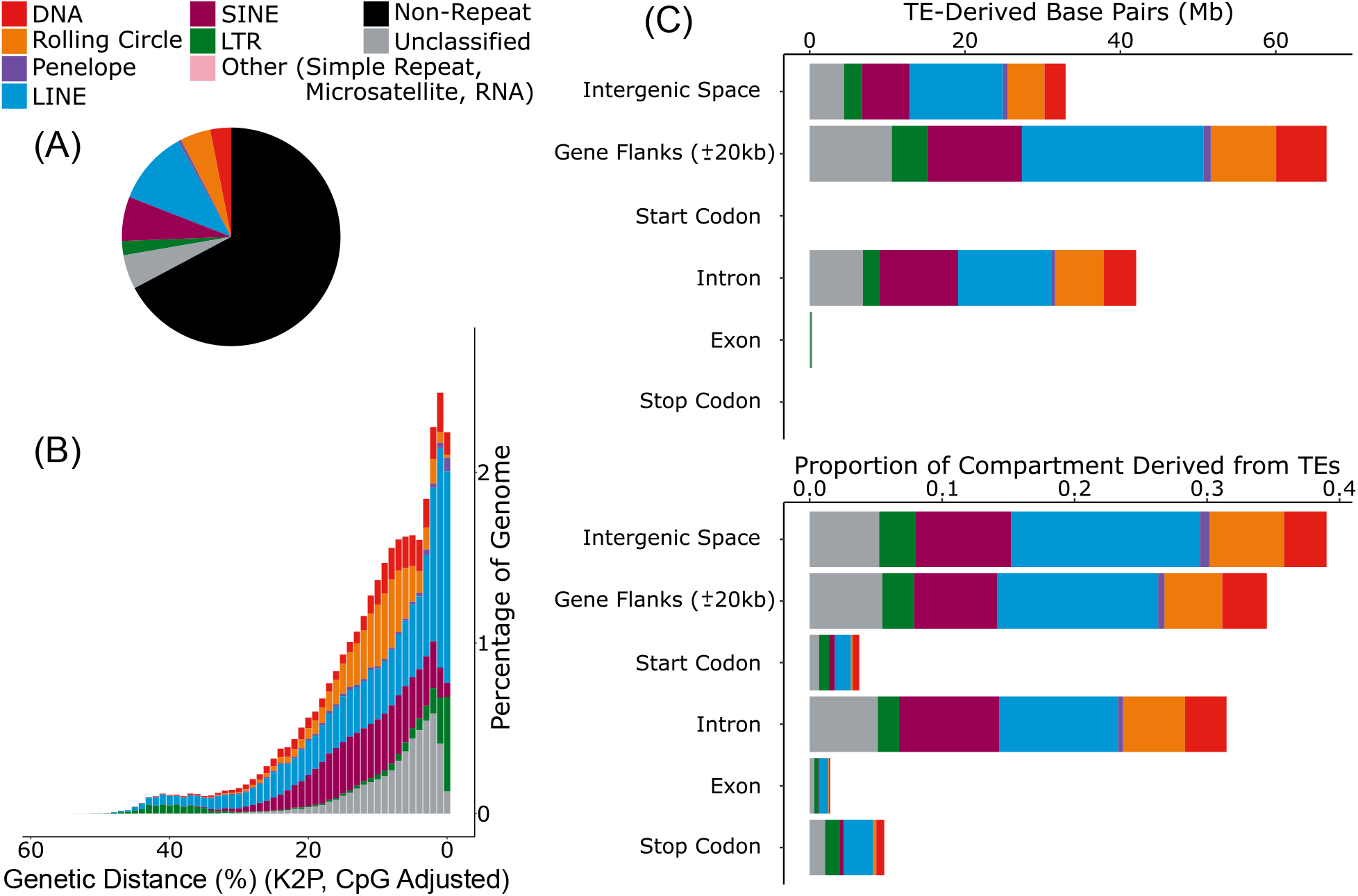
TEs within the genome assembly of *I. podalirius*. (A) The proportion of the assembly comprised of the main TE classifications. (B) A repeat landscape plot illustrating the proportion of repeats in the genome at different genetic distances (%) to their respective RepeatModeler consensus sequence. Genetic distance is calculated under a Kimura two parameter model with correction for CpG site hypermutability. Lower genetic distances suggest more recent activity. (C) The abundance of TEs in different partitions of the genome, shown in bases and as a proportion of the partition.

Considering all TE classifications, most TEs (70.14%) are found in intergenic regions. As expected, TEs are largely absent from exons with only 0.07% of exonic sequence consisting of TEs, likely due to the deleterious effects of TE insertions in exons (Bourque *et al*. 2018; Sultana *et al*. 2017). In contrast, a substantial fraction of intronic sequence (31.47%) is comprised of TEs (Figure 2C). The most abundant TEs in the assembly, LINEs, comprise 14.25% of intergenic space, 12.14% of gene flanks, 8.98% of intronic regions, and 0.69% of exonic regions (Figure 2C).

We annotated the assembly with 17,826 protein coding genes, coding for 20,222 transcripts (1.13 transcripts per gene). At least one Pfam domain was identified along proteins of 9,363 genes (52.5%) and start codons were found in genes coding for 20,163 proteins (99.71%). The median length of genes is 4.0 kb, with the majority (51.7%) consisting of four exons or fewer. The number of gene predictions is higher than in genome annotations for species in the genus Papilio, such as *Papilio dardanus* (12, 795, Timmermans *et al*. 2020) and *P. bianor* (15, 375, Lu *et al*. 2019), but far lower than in the annotation of the *Parnassius Apollo* genome (28, 344, Podsiadlowski *et al*. 2021). Genetic Distance (%) (K2P, CpG Adjusted)

The density of annotated genomic features differs across chromosomes (Figure 3). For example, the proportion of sequence made up of TEs ranges from 24.4% on chromosome 7 to 39.4% on chromosome 29. Similarly, exon density also ranges approximately two-fold across chromosomes, from 3.4% on chromosome 25 to 6.4% on chromosome 30. The density of TEs is negatively correlated with chromosome length (Spearman’s rank correlation, *ρ*_(28)_ = −0.520, *p* = 0.004), whereas exon density and chromosome length are uncorrelated (Figure 3).

**Figure 3:**
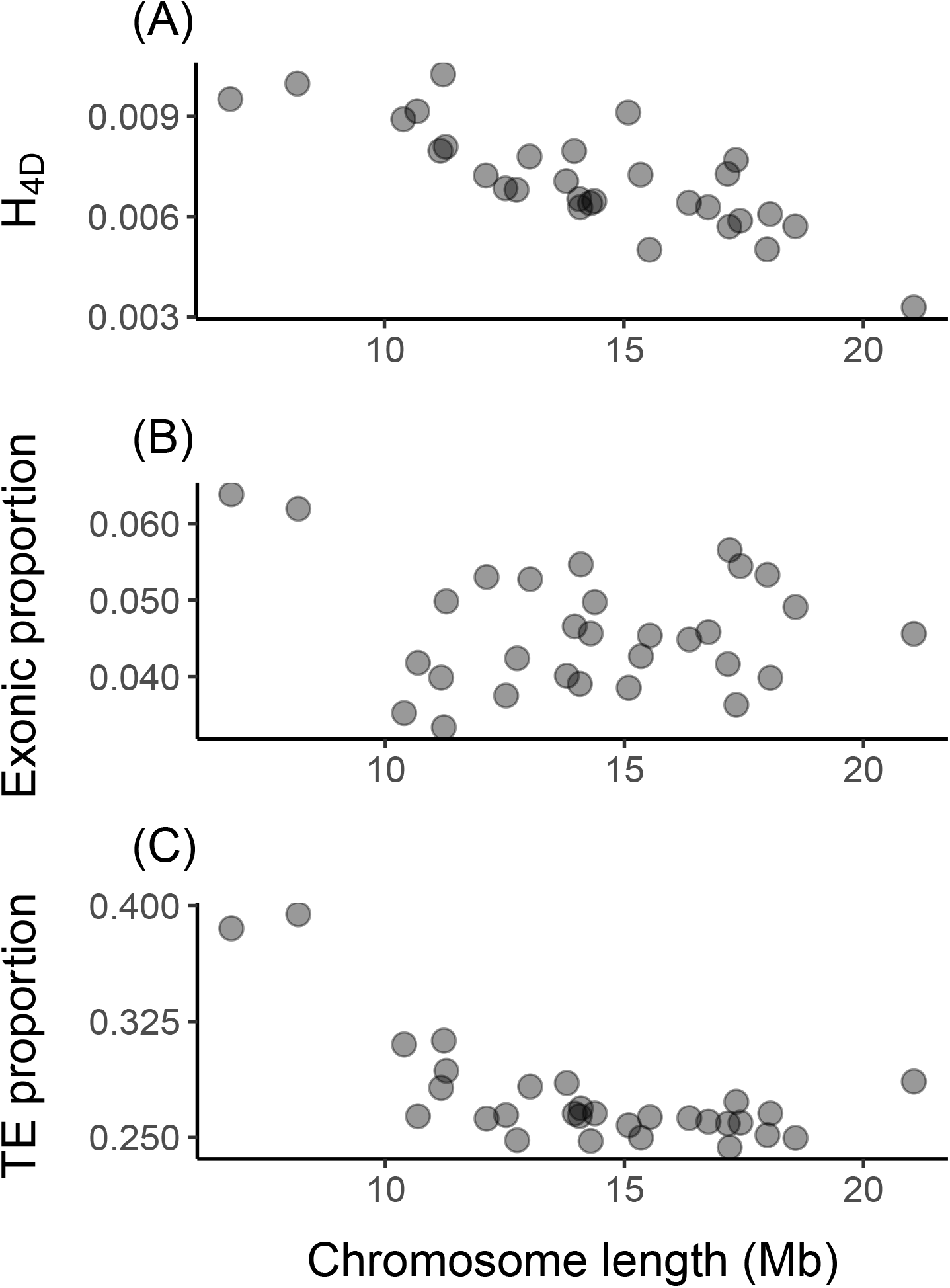
The relationship between chromosome length and (A) heterozygosity at 4D sites, (B) exon density, and (C) TE density.

### 3.3 Genome-wide heterozygosity

Across the genome assembly, we identified 2, 514, 242 heterozygous SNPs in the reference individual MO IP 504, which is equivalent to a per-base heterozygosity (across all sites) of 0.00598 (Table 1). As expected, given selective constraint, heterozygosity in exons is less than half of that in introns and intergenic regions (Table 1). Likewise, within coding sequence heterozygosity is highest for 4D sites and lowest for 0D sites (Table 1), which is expected given that the latter are under strong evolutionary constraint (Sawyer *et al*. 1987).

We note that our estimate of 4D site heterozygosity is comparable, but slightly higher, than *π*_4*D*_ estimates previously reported for *I. podalirius* based on transcriptome assemblies and data from two individuals (0.0052, 0.0057) (Ebdon *et al*. 2021; Mackintosh *et al*. 2019). This difference most likely reflects the fact that transcriptome assemblies are biased towards highly expressed transcripts which experience greater indirect effects of purifying selection (Marek and Tomala 2018).

Heterozygosity at 4D sites (*H*_4*D*_) varies by chromosome: it is lowest on chromosome 1 (0.00328, the putative Z chromosome) and highest on chromosome 25 (0.01026, an autosome). We find a significant negative correlation between chromosome length and *H*_4*D*_ (Spearman’s rank correlation, *ρ*_(28)_ = −0.787, *p* = 2*∗*10^−6^) (Figure 3A).

### 3.4 Conclusions

We describe a chromosome level genome assembly for the scarce swallowtail butterfly *I. podalirius*, with gene and repeat annotation in agreement with previously published butterfly genomes.

When comparing heterozygosity in the reference individual both between chromosomes and between genomic partitions, we recover two well documented patterns of genome-wide sequence variation which result from the direct and indirect effects of selection respectively: i) stark differences in genetic diversity between genomic partitions reflecting differences in selective constraint and ii) a negative correlation between chromosome length and heterozygosity at putatively neutral 4D sites. The latter has been described for several species (including butterflies Martin *et al*. 2016) and is an expected consequence of the fact that the indirect effect of selection on nearby neutral sites depends on the rate of recombination (which tends to be greater for short chromosomes, Haenel *et al*. 2018).

We also find a negative relationship between chromosome length and TE density. This is surprising given that increased recombination rates on shorter chromosomes are expected to result in more efficient selection against TE insertions (Langley *et al*. 1988). Despite this, the observation that smaller chromosomes contain a greater density of repetitive elements has also been reported in *Heliconius* and *Melitaea* butterflies (Cicconardi *et al*. 2021; Ahola *et al*. 2014), suggesting that this may be a conserved feature of Lepidopteran chromosomes.

The *I. podalirius* genome will be valuable resource — not only for genomic analyses that investigate the forces driving genome evolution in the long term — but will also allow for detailed studies of the population level processes that lead to the accumulation of barriers between species still experiencing gene flow.

## 4 Data availability

The genome assembly, gene annotation, and raw sequence data can be found at the European Nucleotide Archive under project accession PRJEB51340. The script used for calculating site degeneracy (partition cds.py), and the script used for visualising HiC contacts (HiC view.py) can be found at the following github repository: https://github.com/A-J-F-Mackintosh/Mackintosh_et_al_2022_Ipod. The mitochondrial genome sequence and the TE annotation can be found at the same repository.

## 5 Acknowledgments

We would like to thank Marian Thompson for preparing the Pacbio sequencing libraries and Andrées de la Filia and Katy MacDonald for help in the molecular lab. We thank Alvaro Martinez Barrio (10X Genomics) and Paul Coupland (Cancer Research UK Cambridge Institute) for help with generating the 10X data. We thank Charlotte Wright for helpful comments on an earlier draft of this manuscript.

