## Supplementary material for "The genome sequence of the scarce swallowtail, *Iphiclides podalirius*"

### 8 Supplementary Materials

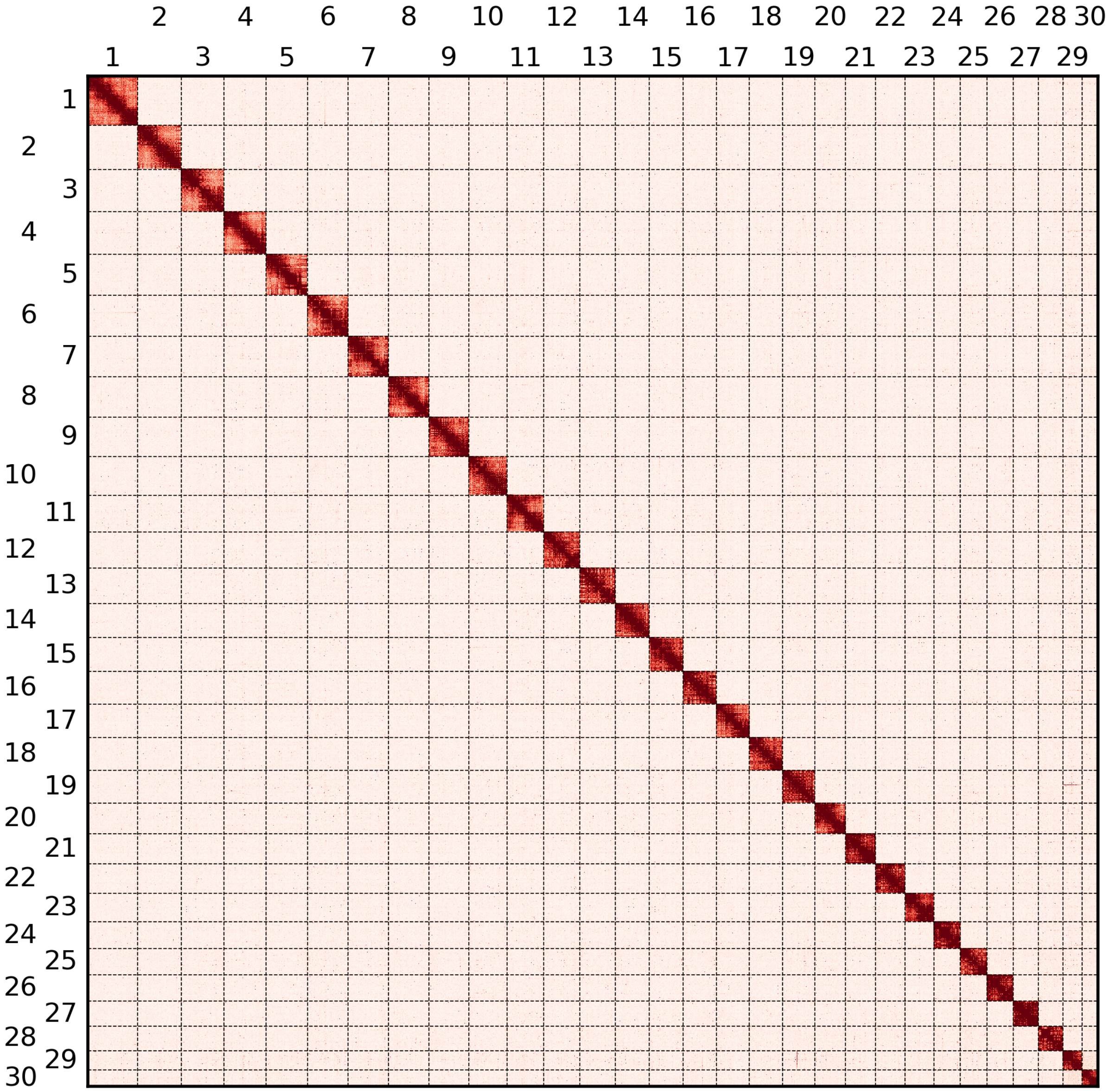


Figure S1: HiC contacts across all 30 *I. podalirius* chromosomes. The intensity of colour is proportional to the number of HiC contacts two regions of the genome share.

Table S2: Annotated transposable elements

| Repeat class | No. elements | Total length (Mb) | Percentage of genome (%) | No. distinct classifications |
|---|---|---|---|---|
| **Retroelement** | 244343 | 86.73 | 20.15 | 1152 |
| SINE | 120173 | 28.19 | 6.55 | 68 |
| LINE | 105889 | 47.40 | 11.01 | 739 |
| Penelope | 6955 | 2.03 | 0.47 | 16 |
| LTR element | 11326 | 9.11 | 2.12 | 329 |
| **DNA transposon** | 39867 | 13.34 | 3.10 | 512 |
| **Rolling Circle** | 81955 | 19.33 | 4.49 | 166 |
| **Unclassified** | 61052 | 21.82 | 5.07 | 355 |
| **Other** | 14 | 0.01 | 0.00 | 2 |
| **Total** | 427241 | 141.23 | 32.81 | 2187 |
